# The Mp*CAFA* gene encodes a ciliary protein required for spermatozoid motility in the liverwort *Marchantia polymorpha*

**DOI:** 10.1101/2025.04.26.650577

**Authors:** Mizuki Morita, Katsuyuki T. Yamato

## Abstract

Bryophytes, pteridophytes, and some gymnosperm species produce motile ciliated spermatozoids that navigate to the egg by regulating ciliary motility in response to a concentration gradient of attractants released from the egg and/or the surrounding cells. However, the structural components of spermatozoid cilia in land plants remain largely unknown. In this study, we investigated MpCAFA (combined calcyphosine [<u>CA</u>PS] with flagellar-associated protein 115 [<u>FA</u>P115]; Mp1g04120) in the liverwort *Marchantia polymorpha*. The N*-*terminal and near C-terminal regions of MpCAFA showed similarity to CAPS, a mammalian EF-hand protein, and FAP115, a ciliary protein of the green alga *Chlamydomonas reinhardtii*, respectively. Mp*CAFA* was expressed specifically in antheridia and its orthologs were found in some algae, bryophytes, pteridophytes, and some gymnosperms, but not in most seed plants. Spermatozoids from mutants lacking functional MpCAFA exhibited a significant decrease in swimming speed. Notably, these mutants showed no obvious morphological defects, including a 9 + 2 axoneme arrangement, and retained chemotactic capability and fertility, forming normal spores. This suggests that MpCAFA is required for spermatozoid motility, but not for sperm chemotaxis or subsequent reproductive processes. The introduction of Mp*CAFA_pro_*:Mp*CAFA*-*mCitrine* fully complemented the mutant phenotype and revealed that MpCAFA-mCitrine was localized along the lengths of the two spermatozoid cilia. Both the CAPS-like and FAP115-like domains were essential for MpCAFA function and subcellular localization in spermatozoid, whereas the C-terminal proline-rich region was dispensable. These findings indicate that MpCAFA is a major ciliary protein in land plants and can serve as a marker for visualizing spermatozoid ciliary movements.

## Introduction

Fertilization is the process by which eggs and sperm fuse to produce offspring. Prior to fertilization, sperm detect specific chemicals released from the egg or its surrounding cells and navigate toward the egg. Navigation is achieved through the precise regulation of ciliary movement in response to the concentration gradient of the chemical attractant. The core structure of the cilium, which is responsible for generating propulsion and controlling the swimming direction, is the axoneme. The axoneme exhibits a conserved 9 + 2 arrangement of microtubules and numerous associated proteins. The structure and function of the axoneme have been intensively studied using animal sperm and the biciliate unicellular green alga *Chlamydomonas reinhardtii*. For example, the sperm of the ascidian *Ciona intestinalis* quickly turn towards the egg upon detection of a decrease in the concentration of attractant released from the egg (Shiba et al. 2008). The mechanism underlying this response involves Ca^2+^ influx triggered by a decrease in attractant levels, followed by the modulation of cilium bending. This modulation is mediated by calaxin, which at high Ca^2+^ concentrations binds to the outer-arm dynein and inhibits microtubule sliding (Mizuno et al. 2012). The intricate structure of microtubule doublets within the *C. reinhardtii* axoneme has been elucidated using single-particle cryo-electron microscopy, revealing how flagella-associated proteins (FAPs) are integrated into the microtubule doublet assembly (Ma et al. 2019).

Motile ciliated spermatozoids are also found in bryophytes, pteridophytes, and some gymnosperms, such as ginkgo and cycads. The number of spermatozoid cilia differs among land plants, ranging from two in bryophytes to as many as 50,000 in the cycad *Zamia pumila* (Norstog 1986). The two cilia of a spermatozoid from liverworts and mosses are arranged in a staggered manner, the anterior and posterior cilia, whereas those from hornworts are oriented side-by-side (Renzaglia and Duckett 1991). In the liverwort *Marchantia polymorpha,* the two cilia exhibit different beat patterns (Miyamura et al. 2018), indicating structural and functional asymmetry between them. Spermiogenesis in *M. polymorpha* is accompanied by an autophagic process that drastically reorganizes cellular components to form a characteristically elongated cell body with two cilia, two mitochondria at its ends, and a plastid at its posterior end (Minamino et al. 2017; Koshimizu et al. 2022; Norizuki et al. 2022). Nevertheless, how cilium asymmetry is established during spermiogenesis, and its biological significance remains to be elucidated.

The core structure of the axoneme, namely the 9 + 2 arrangement, is highly evolutionarily conserved among eukaryotes, including land plants. In addition, an evolutionarily conserved axonemal protein, Parkin co-regulated gene (PACRG) (Dawe et al. 2005), is encoded by the V (male) chromosome and is expressed specifically in the antheridia of *M. polymorpha* (Yamato et al. 2007; Higo et al. 2016), suggesting that PACRG and possibly more proteins are conserved as ciliary proteins in *M. polymorpha* and other land plants with ciliated spermatozoids. Despite the structural conservation of the axoneme, land plants have lost dynein heavy chains α, β, and γ, which are canonical components of axonemes in animals and algae (Lucas and Geisler 2022). In general, two sets of dynein complexes, the outer and inner dynein arms, are attached to each microtubule doublet and generate sliding forces between microtubule doublets, leading to ciliary beating. The outer dynein arm consists of two or three of the heavy chains α, β, and γ (Inaba 2015), and thus the axoneme of land plant spermatozoids lacks outer dynein arms (Hyams and Campbell 1985). In contrast, the cilia of liverworts, hornworts, and some green algae have acquired a unique protein, phosphatidylinositol-binding clathrin assembly protein-K (PICALM-K) (Minamino et al. 2023). PICALM-K is a member of the AP180 N-terminal homology (ANTH) protein family that is highly conserved among eukaryotes, but is distinguished from other members by the presence of a kinase domain at its C-terminal region. PICALM-K is localized in basal bodies and is required for normal cilia function in *M. polymorpha*. These findings suggest that the cilia of land plant spermatozoids have evolved through the loss and gain of their associated proteins.

In our search for novel components of the spermatozoid cilia, we identified and characterized MpCAFA, a novel ciliary protein in *M. polymorpha.* MpCAFA possesses a chimeric structure with similarities between its N*-* and near C-terminal regions with, respectively, mammalian calcyphosine (CAPS), an EF-hand protein, and FAP115, a ciliary protein from *C. reinhardtii*. CAPS is expressed in ciliated cells of humans (Human Protein Atlas; Uhlén et al. 2015), although its function remains unclear. FAP115 binds to the inner surface of the A tubules of the axonemes in *C. reinhardtii* (Ma et al. 2019) and is required for ciliary beating in the protist *Tetrahymena thermophila* (Fabritius et al. 2021). Our results demonstrated that MpCAFA localized to spermatozoid cilia and played a critical role in spermatozoid motility, but did not affect sperm chemotaxis or fertility in *M. polymorpha*. It is expected to serve as a tool for live imaging of cilia when fused with a fluorescent protein.

## Materials and methods

### Plant materials and growth conditions

Male and female accessions of *M. polymorpha*, Takaragaike-1 (Tak-1) and Takaragaike-2 (Tak-2), were used (Chiyoda et al. 2008). Plants were aseptically cultured on 1% (w/v) agar medium containing half-strength Gamborg’s B5 salts (Gamborg et al. 1968), or open cultured on vermiculite soaked with 1/250 Hyponex (Hyponex Japan Co., Osaka, Japan), under continuous fluorescent light (approximately 70 µmol m^-2^ s^-1^) at 23 ± 1°C. Gametangiophores were induced by supplementation with far-red light, as previously described (Chiyoda et al. 2008).

### Preparation of spermatozoids and crossing

Spermatozoids were collected by placing approximately 100 µL of ultrapure water on a mature male receptacle. For crossing, a freshly prepared spermatozoid suspension was applied to female receptacles 2–3 mm in height and diameter, and again 2 days after the initial application. For movie recordings, the spermatozoid suspension collected in a microtube was kept on ice for 5 min, transferred onto a glass slide for 2 min, and then subjected to microscopic observation.

### Quantitative analysis of spermatozoid motility

Spermatozoid movement was observed under a phase-contrast microscope (BX48; Evident, Tokyo, Japan) and recorded at a resolution of 1,280 × 1,024 pixels, a rate of 100 frames per second (fps), and a shutter speed of 1/300 s using a high-speed digital camera (HAS-U1; Ditect, Tokyo, Japan). Spermatozoids were tracked using TrackMate ver. 7.11.1 (Tinevez et al. 2017), a plugin bundled with the Fiji distribution of ImageJ, with the following modified parameters: estimated blob diameter, 30.606 pixels; initial threshold, 0.2; linking max distance, 30.606 pixels; gap-closing max distance, 30.606 pixels; and gap-closing max frame gap, 2. The trajectory of each spermatozoid was generated by applying moving averages to its raw coordinates, with a window size of 20. The linearity of each spermatoid trajectory was calculated as the ratio of the displacement (linear distance between the start and end points of the trajectory) to the trajectory.

### Promoter-GUS assay

The 3,084-bp Mp*CAFA* promoter region, including the 5’-UTR and 6 bp of coding sequence, Mp*CAFA_pro_*, was amplified with primers (Table S1) from Tak-1 genomic DNA by PCR and subcloned into the pENTR/D-TOPO vector (Thermo Fisher Scientific, Waltham, MA, USA). The inserted fragment was incorporated into pMpGWB104 using Gateway LR Clonase II Enzyme mix (Thermo Fisher Scientific; Fig. S1). To generate transgenic plants, these binary vectors were introduced into germinating spores (Ishizaki et al. 2008) using *Agrobacterium tumefaciens* GV2260. Plants were soaked in β-glucuronidase (GUS) staining solution (5.7 mM X-Gluc [5-bromo-4-chloro-3-indolyl-β-glucuronide]; Bio Medical Science Inc., Tokyo, Japan), 1.5 mM K_3_Fe(CN)_6_ (Nacalai Tesque, Kyoto, Japan), 1.5 mM K_4_Fe(CN)_6_ (Nacalai Tesque), and 0.9% Triton X-100 (Nacalai Tesque) and incubated at 37℃ overnight.

### Generation of MpCAFA-knockout and complementation lines

The Mp*cafa^ge^* lines were generated using the CRISPR/Cas9 protocol for *M. polymorpha* (Sugano et al. 2018). Two guide RNAs (gRNAs) were designed to target coding sequences in the third and fifth exons at sites 1 and 2, respectively (Table S1). Double-stranded DNA for each gRNA was generated by annealing complementary oligonucleotides and ligated with the entry vector pMpGE_En03 digested with BsaI using DNA Ligation Kit Mighty Mix (Takara Bio Inc., Kusatsu, Japan). The gRNA expression cassette for each entry vector was transferred to the destination binary vector pMpGE010 using Gateway LR Clonase II Enzyme mix (Thermo Fisher Scientific). To generate transgenic plants, germinating spores were infected with *A. tumefaciens* GV2260 carrying binary vectors (Ishizaki et al. 2008).

For complementation, the Mp*CAFA* gene, including its promoter region, was amplified from Tak-1 genomic DNA by PCR with the primers listed in Table S1 and subcloned into the entry vector pENTR/D-TOPO (Thermo Fisher Scientific). The Mp*CAFA* gene was then transferred to the destination binary vector pMpGWB347 (Ishizaki et al. 2015) using the Gateway LR Clonase II Enzyme mix (Thermo Fisher Scientific; Fig. S1). A Mp*cafa^ge^* plant without the CRISPR/Cas9 construct was selected from the offspring of Tak-2 × Mp*cafa^ge^*-1 #9-2 (Mp*cafa^ge^*-1 #9-2_4). To generate transgenic plants, gemmae of Mp*cafa^ge^*-1 #9-2_4 were infected with *A. tumefaciens* GV2260 carrying a binary vector (Tsuboyama-Tanaka and Kodama 2015).

### Generation of transgenic lines expressing the MpCAFA domains separately

The coding sequences of the CAPS, FAP115, and FAP115+P-rich domains of Mp*CAFA* were amplified from Tak-1 genomic DNA by PCR with the primers listed in Table S1, and inserted downstream of Mp*CAFA_pro_* in the entry vector constructed for the promoter-GUS assay described above. The expression cassettes were transferred to the pMpGWB101, pMpGWB301, pMpGWB129, and pMpGWB347 destination binary vectors (Ishizaki et al. 2015) using Gateway LR Clonase II Enzyme mix (Thermo Fisher Scientific; Fig. S1). To generate transgenic plants, gemmae of Mp*cafa^ge^*-1 #9-2_4 were infected with *A. tumefaciens* GV2260 carrying the binary vectors (Tsuboyama-Tanaka and Kodama 2015).

### Fluorescent microscopy

Mature antheridia were isolated from antheridiophores and observed under a fluorescence stereomicroscope (MVX10; Evident) equipped with an sCMOS camera (Zyla 4.2 PLUS; Andor, Belfast, UK). Spermatozoids were stained with Kakshine PC3 (Uno et al. 2021) to visualize the nuclei and were observed under an all-in-one fluorescence microscope (BZ-X810; KEYENCE, Osaka, Japan).

### Electron microscopy

For the observation of spermatozoid cilia using transmission electron microscopy, mature antheridia were pre-fixed in 50 mM cacodylate buffer (pH 7.4) containing 2% (v/v) paraformaldehyde and 2% (v/v) glutaraldehyde at 4°C overnight. The subsequent processes were performed at Tokai Electron Microscopy, Inc. (Nagoya, Japan). The prefixed antheridia were washed three times in 50 mM cacodylate buffer at 4°C for 30 min, and then postfixed in 50 mM cacodylate buffer containing 2% (w/v) osmium tetroxide at 4°C for 3 h. The fixed antheridia were dehydrated in an ascending series of ethanol concentrations as follows: 50% ethanol at 4°C for 30 min, 70% ethanol at 4°C 30 min, 90% at room temperature for 30 min, and 100% at room temperature for 30 min (three times) and overnight. The dehydrated antheridia were then equilibrated with propylene oxide (PO) at room temperature for 30 min (twice) and with a 1:1 mixture of PO and Quetol-651 (Nisshin EM Co., Tokyo, Japan) at room temperature for 3 h. They were then embedded in Quetol-651 at 60°C for 48 h. Ultrathin sections (80-nm thick) were cut with a diamond knife using an ultramicrotome (Leica Ultracut UCT, Vienna, Austria), mounted on copper grids, and stained with 2% (w/v) uranyl acetate at room temperature for 15 min, and then with lead stain solution (Sigma-Aldrich, St. Louis, MO, USA) at room temperature for 3 min. The stained grids were imaged using a transmission electron microscope (JEM-1400Plus; JEOL Ltd., Tokyo, Japan) operating at 100 kV and equipped with a CCD camera (EM-14830RUBY2, JEOL Ltd.).

### Phylogenetic analysis

The amino acid sequences of MpCAFA and its homologs were collected from public databases (Table S2) and aligned to construct phylogenetic trees using the MUSCLE algorithm implemented in the phylogenetic analysis software MEGA 11 (Kumar et al. 2018; Stecher et al. 2020). CAPS-like and FAP115-like domains were used separately to construct phylogenetic trees for CAPS and FAP115 homologs, respectively.

## Results

### CAFA is conserved among streptophytes having motile spermatozoids or ciliated cells

To identify novel components of spermatozoid cilia, we searched for genes predominantly or specifically expressed in antheridium using the *Marchantia* reference genome ver. 3.1 (Bowman et al. 2017) and RNA-seq data (Higo et al. 2016), and identified Mapoly0005s0195 (currently Mp1g04120 in the genome version 7.1), which encodes a protein with an EF-hand Ca^2+^-binding domain. The N- and C-terminal regions of the protein were similar to those of <u>CA</u>PS, a human EF-hand protein (47%), and <u>FA</u>P115, a ciliary protein of the green alga *C. reinhardtii* (49%), respectively (Fig. 1); thus, we referred to it as MpCAFA. The FAP115-like domain was followed by a proline-rich domain, which was found only in *M. polymorpha*, *M. paleacea*, and *Riccia fluitans* among the species investigated in this study, implying that the domain was acquired in the Marchantiales lineage (Table 1). Human CAPS, its closely related proteins calcyphosine 2 (CAPS2) and calcyphosine-like, and *C. reinhardtii* FAP115 have multiple conserved EF-hands, but only one was detected within the CAPS-like domain of MpCAFA (represented by asterisks in Fig. 1). Phylogenetic analysis revealed that the CAPS domain of MpCAFA belonged to the CAPS2 clade, and its homologs were not detected in angiosperm genomes (Fig. 2a). FAP115 was conserved in ciliates, green algae, and land plans with ciliated cells (Fig. 2b).

**Fig. 1.**
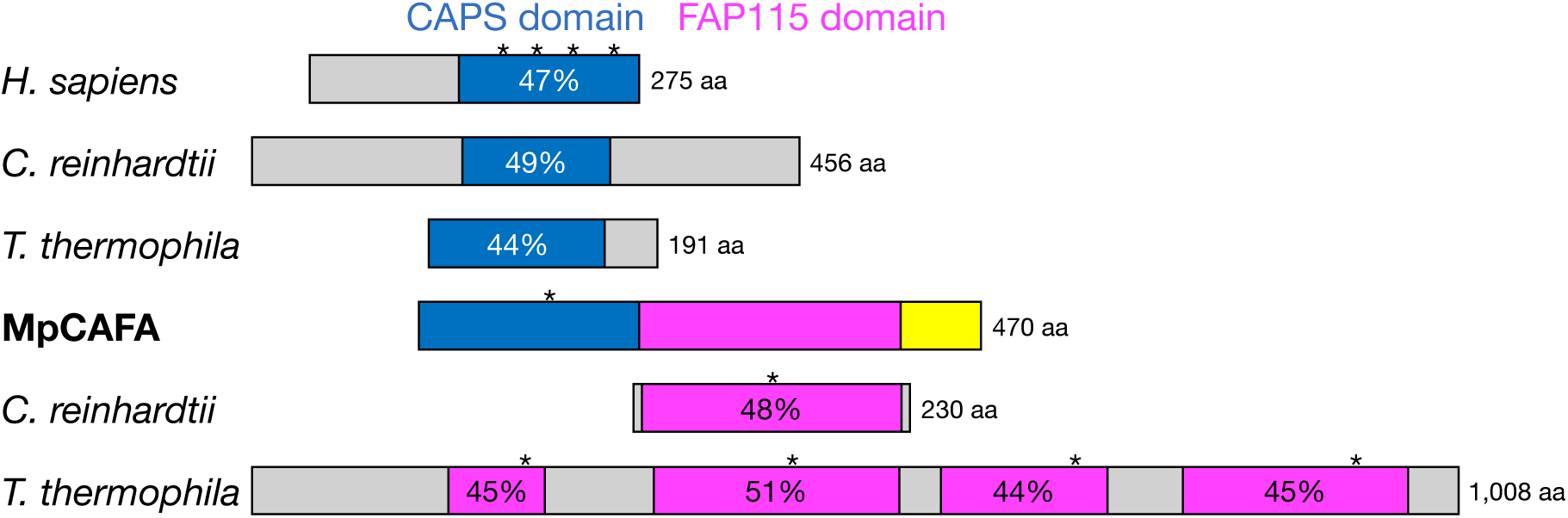
MpCAFA consisted of conserved CAPS- and FAP115-like domains. Schematic illustration of the structure of MpCAFA and comparison with their homologs. The CAPS-like, FAP115-like, and proline-rich domains are shown in blue, magenta, and yellow, respectively. Asterisks indicate the locations of predicted EF-hands. The percentages in the domains of other species indicate the similarity to those of MpCAFA. The accession numbers of these sequences are listed in Table S2.

**Table 1.**
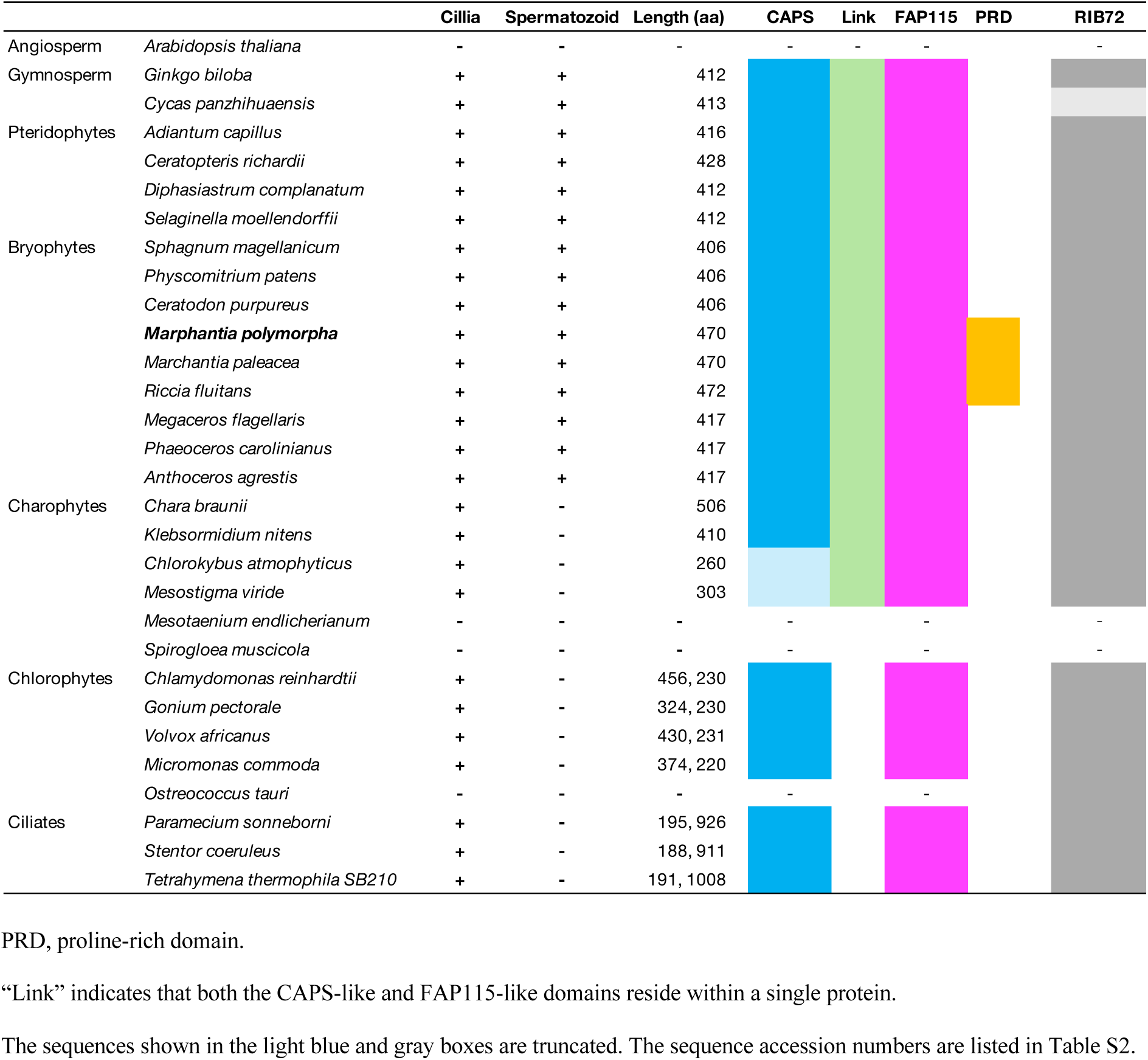
Distribution of cilia and CAFA-related proteins in Viridiplantae and ciliates

**Fig. 2.**
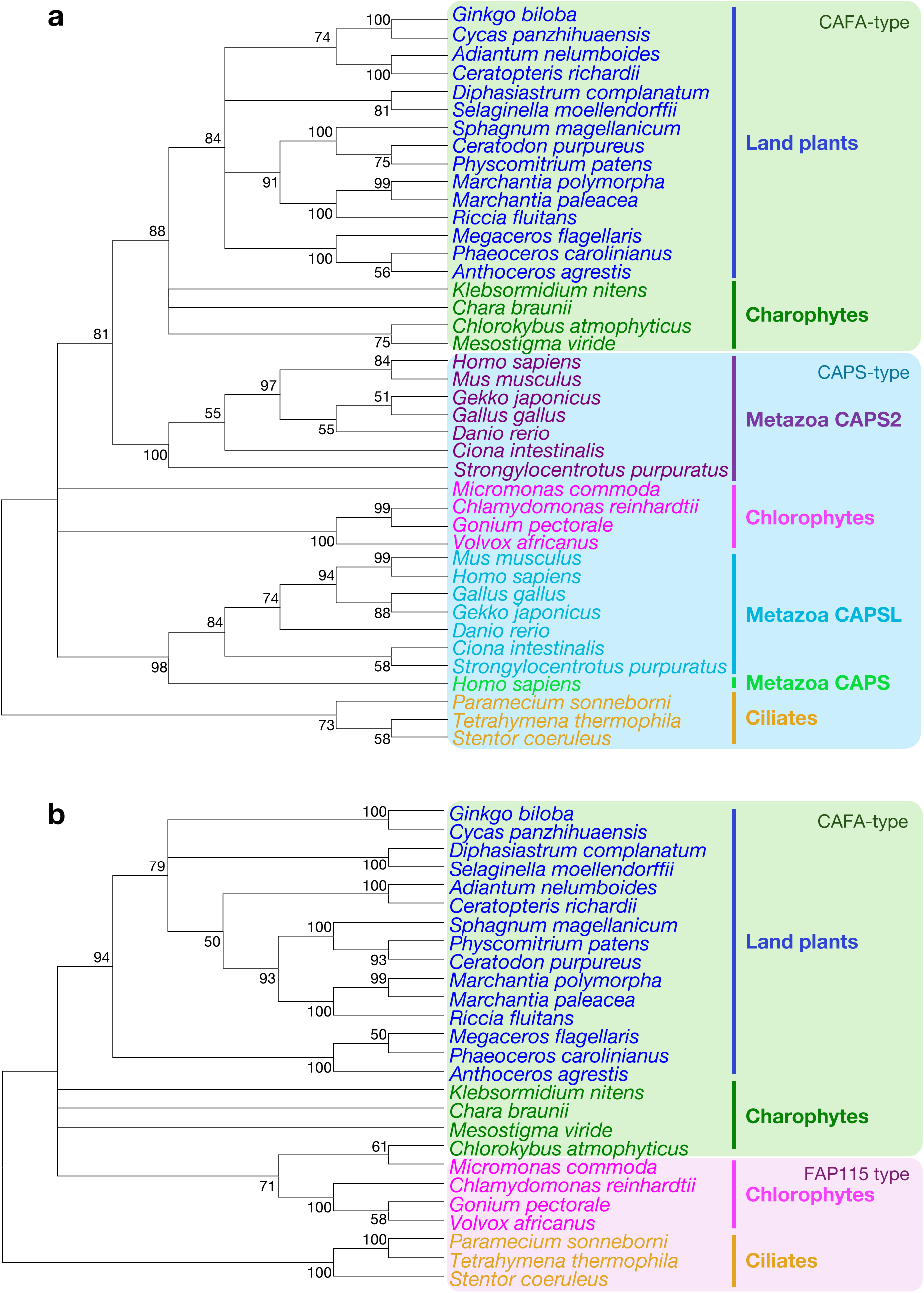
Phylogenetic analyses of conserved CAPS- and FAP115-like domains. Phylogenetic trees of the CAPS-like (**a**) and FAP115-like (**b**) domains. Phylogenetic trees were constructed using the neighbor-joining method with the Jones-Taylor-Thornton model. The numbers at the nodes indicate the support values obtained with 1,000 bootstrap replicates using MEGA11 software.

Homologs of MpCAFA, defined by the contiguous presence of both the CAPS and FAP115 domains, were found only in bryophytes, pteridophytes, cycads, ginkgo and charophytes, excluding conjugating green algae (Zygnematophyceae)(Table 1; Figs. 2 and 3). The concurrent absence of CAFA and cilia in Zygnematophyceae suggests a potential functional link between CAFA and ciliary development and movement. Conversely, ciliated Chlorophyta, such as *C. reinhardtii* and some ciliates, possess separate CAPS- and FAP115-like proteins (Table 1), reinforcing the hypothesis that these domains are associated with ciliary functions.

**Fig. 3.**
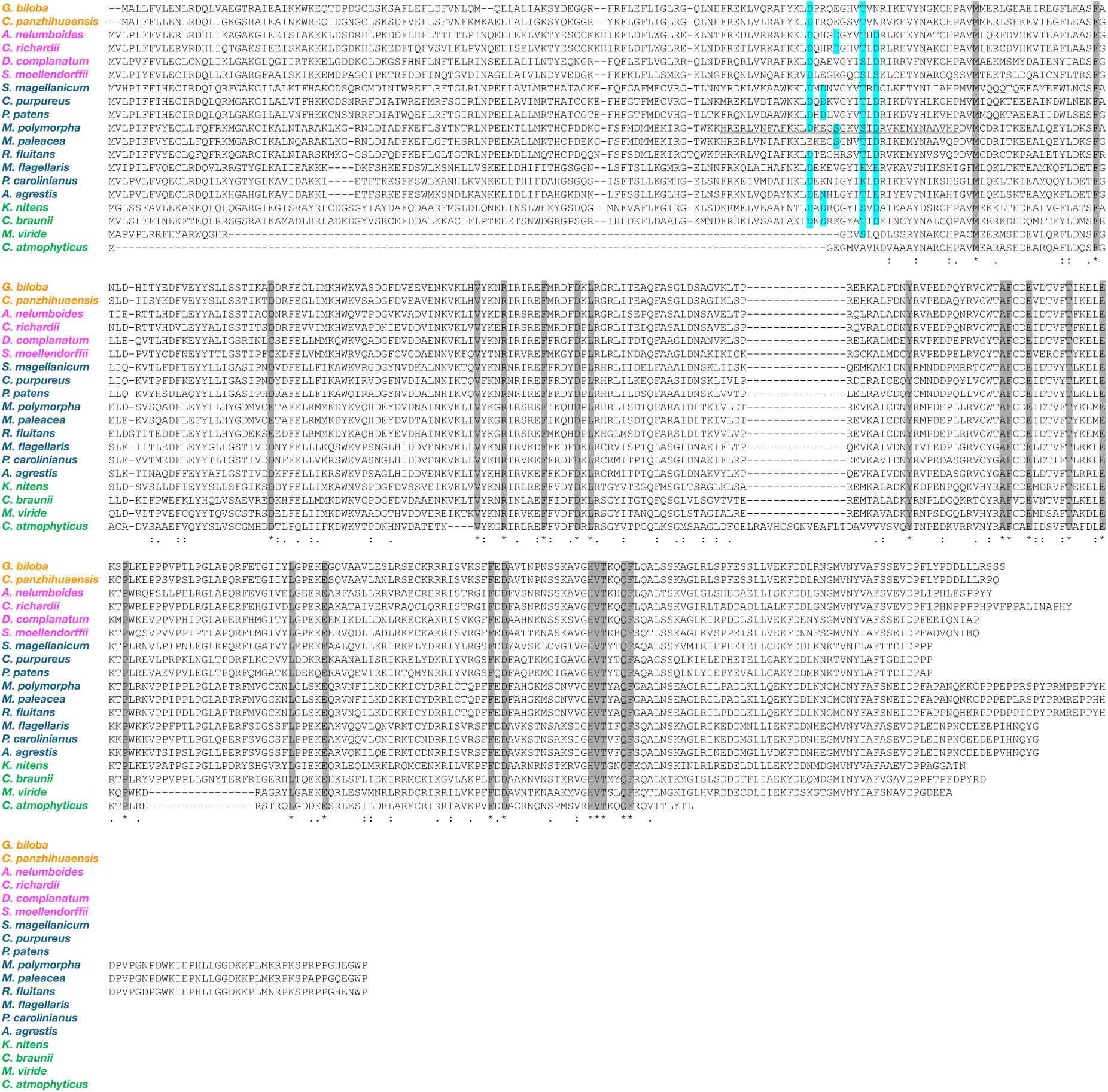
Multiple sequence alignment of MpCAFA and its homologs. Gymnosperm, pteridophytes, bryophytes, and charophytes are shown in orange, magenta, blue, and green, respectively. The EF-hand domain predicted by InterProScan in *M. polymorpha* is underlined and the amino acid residues predicted to be involved in calcium binding are highlighted in cyan. Amino acid residues conserved among the species shown here are highlighted in gray.

### MpCAFA is specifically expressed in antheridia

Mp*CAFA* was identified *in silico* as one of the genes predominantly expressed in antheridia. To verify its promoter activity *in planta*, transgenic lines expressing the *GUS* gene under the control of the Mp*CAFA* promoter (Mp*CAFA_pro_*:*GUS*) were generated. Consistent with publicly available RNA-seq data (MarpolBase Expression; https://mbex.marchantia.info/chroexp/Mp1g04120/), GUS activity was specifically detected in antheridia (Fig. 4).

**Fig. 4.**
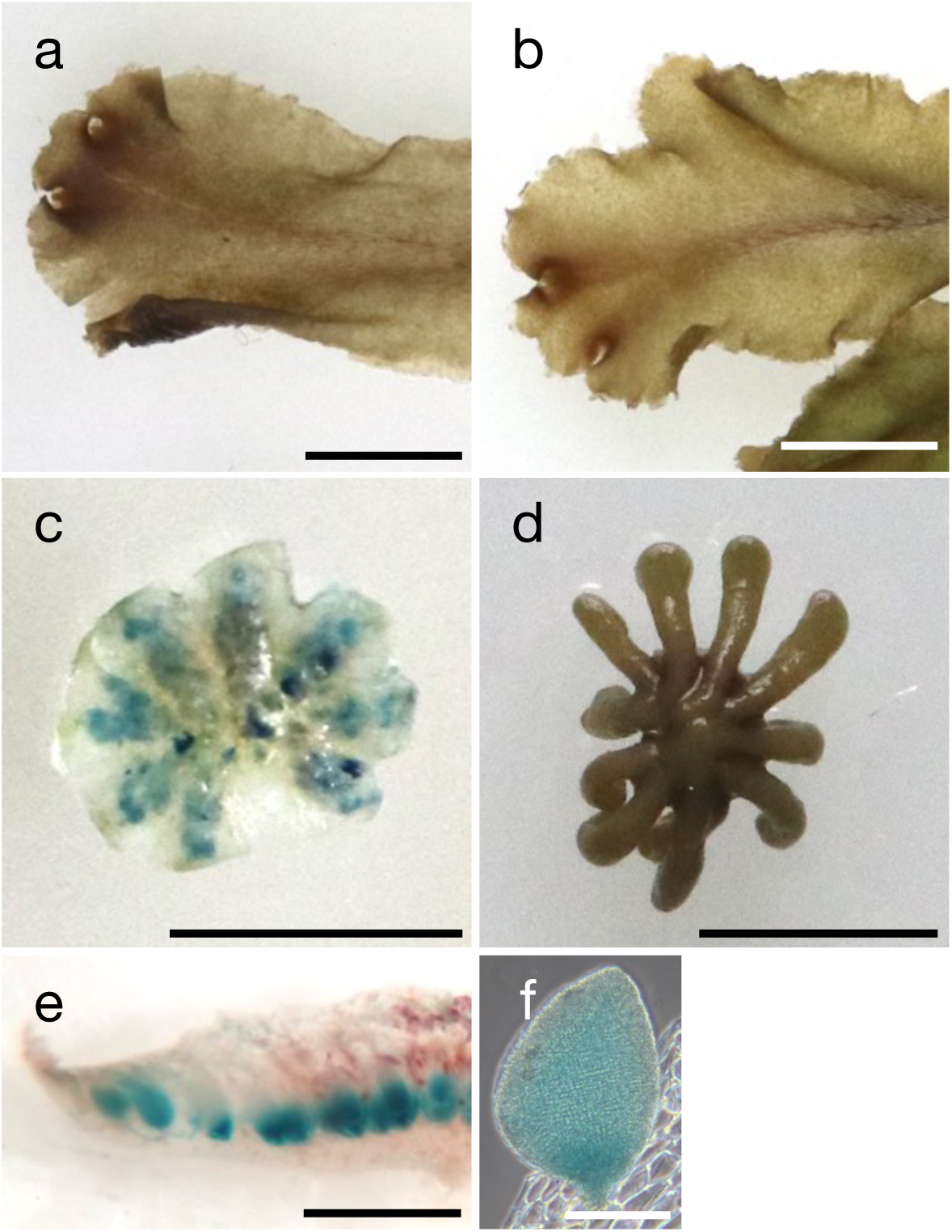
Antheridium-specific expression of Mp*CAFA*. GUS staining of plants carrying the Mp*CAFA_pro_*:*GUS* construct. Male (**a**) and female (**b**) thalli, antheridiophore (**c**), archegoniophore (**d**), longitudinal sections of male receptacle (**e**), and antheridium (**f**) are shown. Scale bars: 1 cm (**a** and **b**), 5 mm (**c** and **d**), 1 mm (**e**), and 250 µm (**f**).

### MpCAFA is required for proper spermatozoid motility, but not for fertility

To characterize the function of MpCAFA, loss-of-function mutants Mp*cafa*^ge^, were generated by CRISPR/Cas9-mediated genome editing (Sugano et al. 2018). Two target sites within the CAPS-like domain were selected (Fig. 5a) and three independent Mp*cafa*^ge^ plants were obtained for each site and sex (Figs. 5b, c and S2). None of the Mp*cafa*^ge^ plants exhibited obvious growth or morphological defects in the thalli or receptacles (Fig. S3). Given the antheridium-specific expression of Mp*CAFA*, suggesting its role in spermatozoid development and/or function, we investigated the spermatozoids of Mp*cafa*^ge^ plants. Morphologically normal spermatozoids were released from all male Mp*cafa*^ge^ plants upon the application of water to their receptacles (Fig. 6a, b), and the cilia of spermatozoids from Mp*cafa*^ge^ retained the typical 9 + 2 arrangement (Fig. 6c, d), indicating that MpCAFA is not essential for normal spermatozoid development. However, most Mp*cafa*^ge^ spermatozoids displayed reduced motility compared to the wild-type spermatozoids, with some exhibiting aberrant swimming behaviors, such as circling (Movies S1–S3). To further investigate the motility defect, the trajectories of individual wild-type and Mp*cafa*^ge^ spermatozoids were tracked and analyzed (Fig. 7a). Although the swimming direction of most wild-type spermatozoids was predominantly straight, Mp*cafa*^ge^ spermatozoids frequently exhibited circular trajectories or increased swing widths (Fig. 7a). We examined two aspects of spermatozoid swimming behavior: swimming speed and linearity (see Materials and Methods). The average swimming speed of spermatozoids significantly decreased in Mp*cafa*^ge^ plants, and a group of Mp*cafa*^ge^ spermatozoids showed particularly low motility, which was not observed in wild-type spermatozoids (Fig. 7b). Mp*cafa*^ge^ spermatozoids also showed less straight swimming trajectories than wild-type spermatozoids (Fig. 7c). These observations suggested that MpCAFA was required for proper ciliary beating to generate propulsion and to orient swimming direction.

**Fig. 5.**
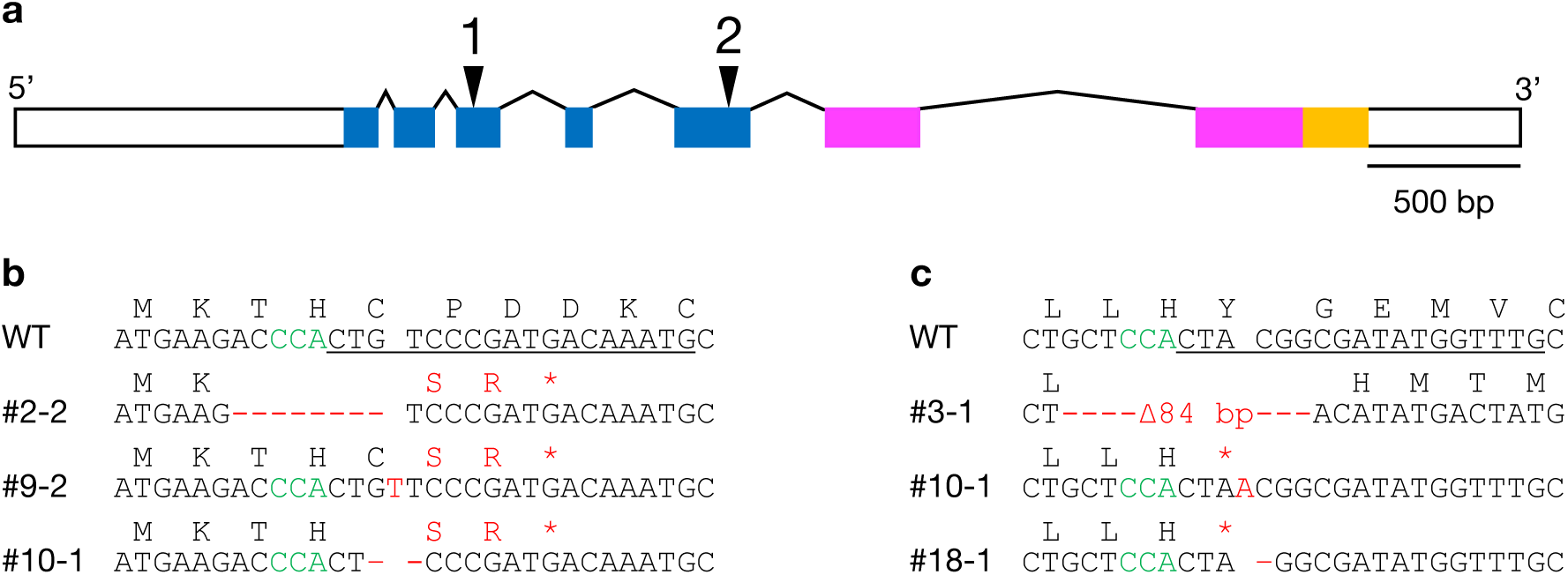
Generation of male loss-of-function Mp*cafa^ge^* mutants. (**a**) Schematic illustration of the Mp*CAFA* gene. Open and filled boxes indicate untranslated and coding regions, respectively. CAPS- and FAP115-like domains and the proline-rich domain are shown in blue, magenta and orange, respectively. Triangles indicate the target sites for CRISPR/Cas9. (**b**) and (**c**) Mutations introduced by CRISPR/Cas9 at the target sites 1 and 2 in (**a**), respectively. Nucleotide sequences of the target sites in the wild-type (WT) and male mutant (*e.g.*, #2-2) plants are shown with the translated amino acid sequence above for each. The target and protospacer adjacent motif (PAM) sequences are underlined and in green, respectively. Mutations are shown in red. The large deletion is indicated with the number of deleted nucleotides. Asterisks indicate translational stops introduced by genome-editing. Sequences in female mutants are given in Fig. S2.

**Fig. 6.**
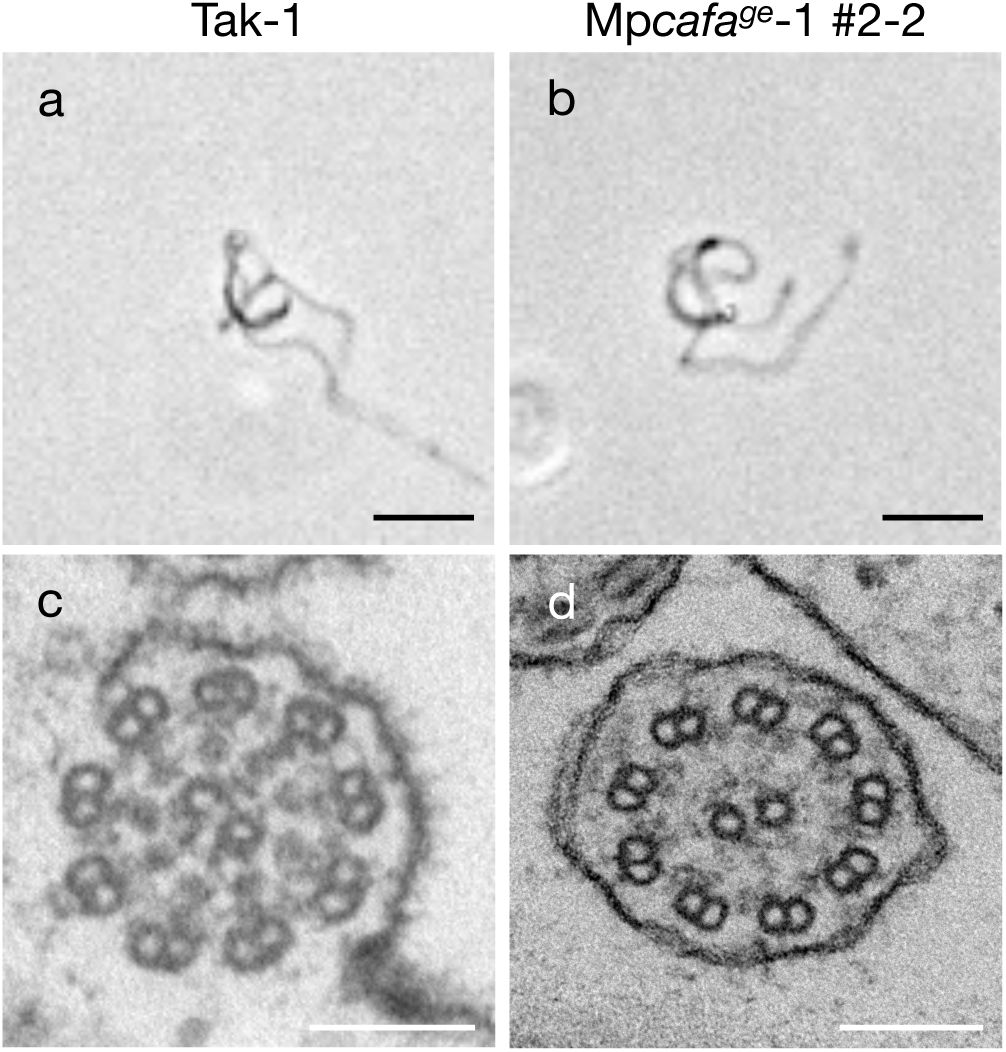
Morphology and axoneme structure were normal in Mp*cafa^ge^* spermatozoids. (**a**) and (**b**) Phase-contrast images of spermatozoids from wild-type male Takaragaike-1 (Tak-1) and representative Mp*cafa^ge^*-1 #2-2, respectively. Scale bar, 10 µm. (**c**) and (**d**) transmission electron microscope images of the cross-sections of cilia from Tak-1 and Mp*cafa^ge^* #2-2 spermatozoids, respectively. Scale bar, 100 nm.

**Fig. 7.**
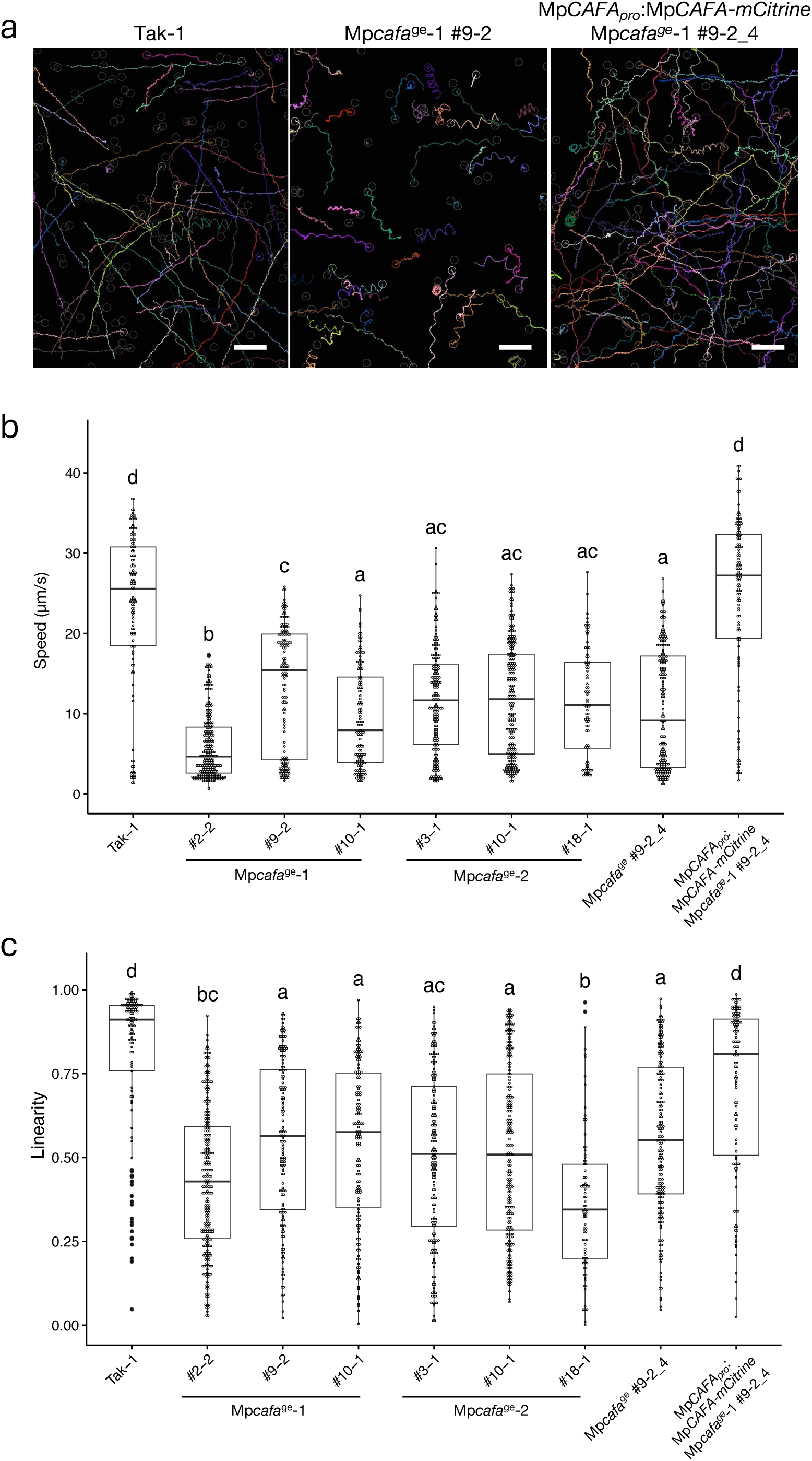
Spermatozoid motility was impaired in Mp*cafa^ge^* plants. (**a**) Trajectories of wild-type (Tak-1), representative mutant (Mp*cafa^ge^*-1 #9-2), and complemented (Mp*CAFA_pro_*:Mp*CAFA*-*mCitrine* Mp*cafa^ge^*-1 #9-2_4) spermatozoids. Mp*cafa^ge^*-1 #9-2_4 is one of the progenies obtained by crossing Mp*cafa^ge^* #9-2 with a wild-type female and it lacks the CRISPR/Cas9 construct. Each spermatozoid was tracked for 10 seconds. Scale bar, 50 µm. (**b**) and (**c**) Box-and-whisker plots of spermatozoid swimming speed and linearity, respectively. The upper and lower limits indicate the first and third interquartile ranges, respectively, and the middle bars indicate the medians. Whiskers indicate ± 1.5× the interquartile ranges. Different lowercase letters indicate significant differences (*P* < 0.05; Tukey–Kramer test). n = 121, 245, 149, 139, 177, 223, 83, 188, and 114 spermatozoids of Tak-1; Mp*cafa*^ge^-1 #2-2, #9-2 and #10-1; Mp*cafa*^ge^-2 #3-1, #10-1, and #18-1; Mp*cafa*^ge^-1 #9-2_4, and Mp*CAFA_pro_*:Mp*CAFA*-*mCitrine* Mp*cafa^ge^*-1 #9-2_4, respectively.

Since it is essential for spermatozoids to navigate to eggs to achieve fertilization, we investigated whether MpCAFA is required for the chemotactic behavior of spermatozoids. Wild-type archegonia were excised from a female receptacle and placed with wild-type or Mp*cafa^ge^* spermatozoids in water. Under natural conditions, spermatozoids are attracted by unidentified substances released from the opening at the end of an archegonium, after which, they enter and swim through the neck to reach an egg that resides at the base of the archegonium, which is most likely the source of the spermatozoid attractant. However, when an archegonium excised from a female receptacle was used, spermatozoids were attracted by its base far more strongly than by the tip of its neck (Fig. 8, top), likely because the attractant was released from the wound at the base. Mp*cafa^ge^* spermatozoids exhibited impaired motility, but were strongly attracted to archegonia (Fig. 8, top), suggesting that MpCAFA is dispensable for spermatozoid chemotactic behavior. Next, we tested whether Mp*cafa^ge^* spermatozoids were fertile by applying them to wild-type female archegoniophores. All female plants crossed with spermatozoids from Mp*cafa^ge^* lines produced spores (Fig. 8, bottom), indicating that MpCAFA is not essential for fertility.

**Fig. 8.**
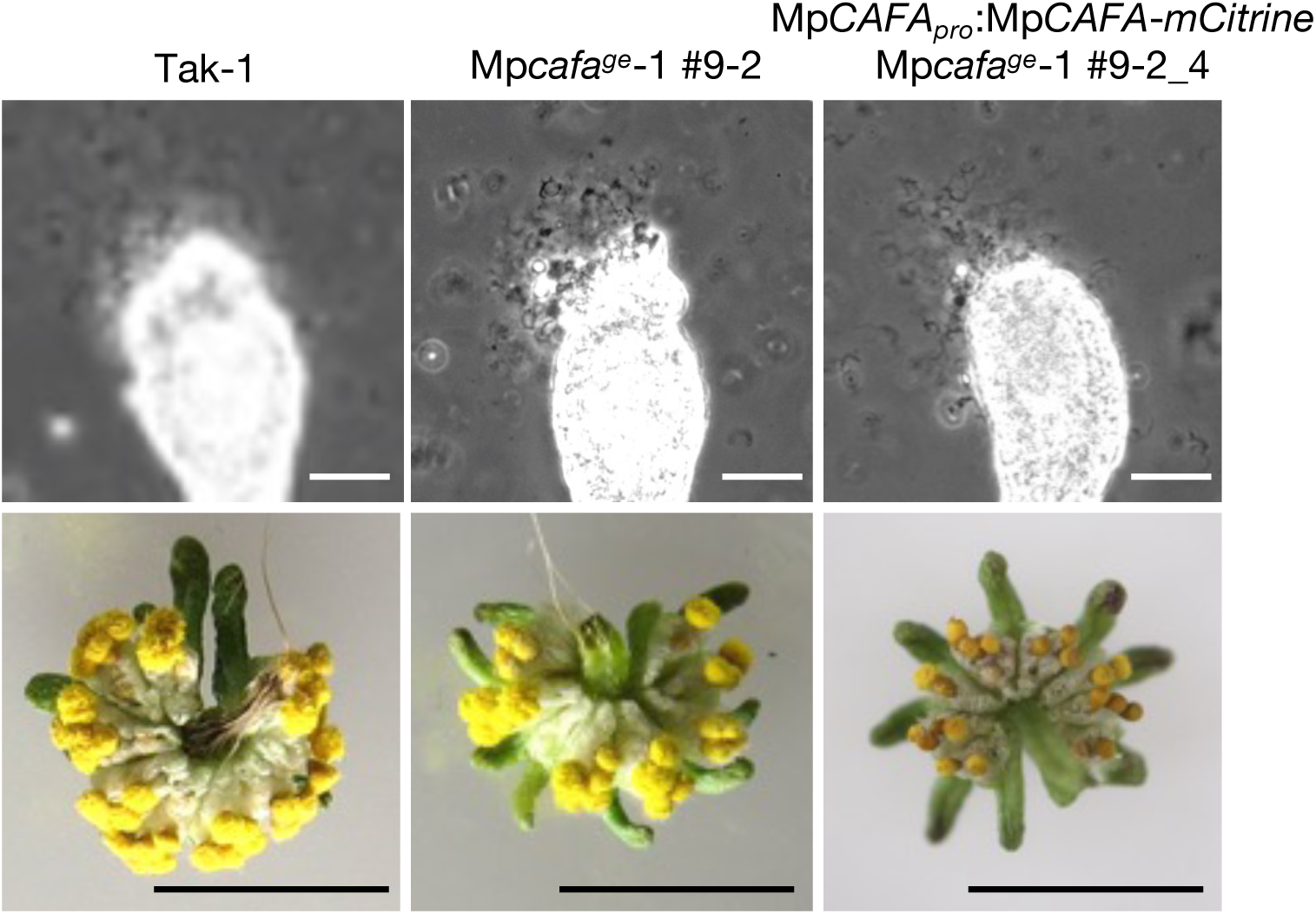
Mp*cafa^ge^* spermatozoids were attracted to an archegonium and were fertile. **Top**: Attraction of wild-type, Mp*cafa^ge^*, and Mp*cafa^ge^*-1 #9-2, Mp*CAFA_pro_*:Mp*CAFA*-*mCitrine* Mp*cafa^ge^*-1 #9-2_4 spermatozoids, to wild-type archegonia. Spermatozoids were attracted to the base of each archegonium, instead of the tip of the neck. Scale bar, 30 µm. **Bottom**: Spores formed by crossing wild-type, Mp*cafa^ge^*, and Mp*CAFA_pro_*:Mp*CAFA*-*mCitrine* Mp*cafa^ge^*-1 #9-2_4 spermatozoids with wild-type females. Scale bar, 1 mm.

These results indicated that MpCAFA is not essential for chemotaxis, fertilization, sporophyte development, and sporogenesis, but is essential for spermatozoid motility, likely for ciliary functionality.

### MpCAFA is localized along the entire length of both spermatozoid cilia

To investigate the subcellular localization of the MpCAFA protein in spermatozoid, MpCAFA was fused with mCitrine at its C-terminus (Mp*CAFA*_pro_:Mp*CAFA-mCitrine*) and the construct was introduced into Mp*cafa^ge^* plants. Spermatozoids from the obtained plants showed swimming speeds and linearity similar to those of Tak-1 (Fig. 7; Movie S5), indicating that Mp*CAFA*_pro_:Mp*CAFA-mCitrine* functionally complemented the mutant phenotype of Mp*cafa^ge^.* The mCitrine signal was detected in antheridia at all developmental stages (Fig. 9a) and along the entire length of both cilia of spermatozoids from plants complemented with Mp*CAFA*_pro_:Mp*CAFA-mCitrine* (Fig. 9b), indicating that MpCAFA is a ciliary component. The mCitrine signal at the basal ends of the two cilia revealed a staggered arrangement of the anterior and posterior cilia (Fig. 9c), indicating that MpCAFA was localized in the axoneme.

**Fig. 9.**
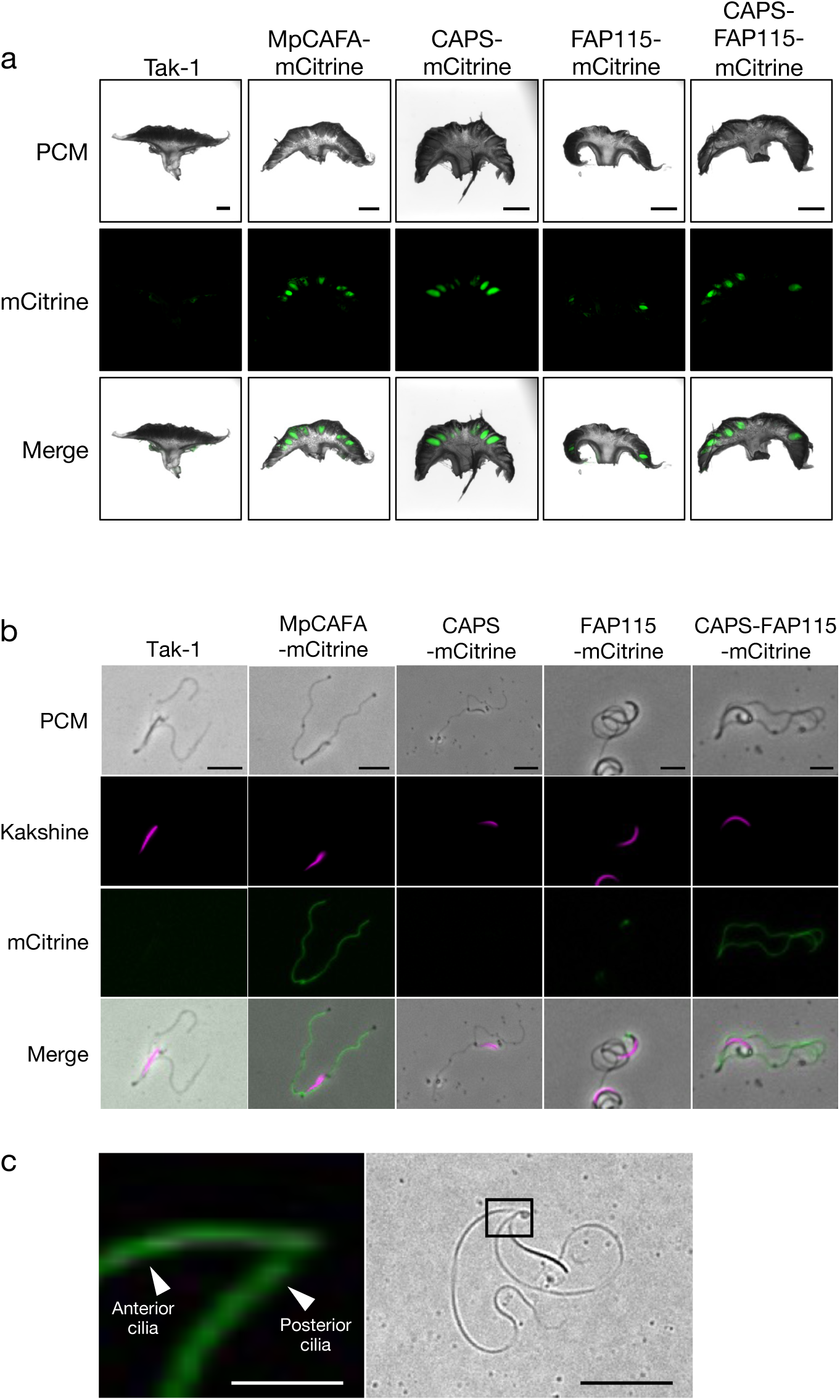
Localization of MpCAFA and its truncated proteins. (**a**) and (**b**) Antheridiophores and spermatozoids from Mp*cafa^ge^*-1 #9-2_4 plants expressing the Mp*CAFA*-promoter-driven constructs shown above. PCM, phase-contrast microscopy. Scale bar, 10 µm. (**c**) The basal ends of cilia expressing MpCAFA-mCitrine (left) are shown enlarged from the boxed area in the right panel. Scale bars, 25 and 100 µm in the left and right panels, respectively. **c** Expression of mCitrine-tagged proteins in male receptacles. Scale bar, 1 mm.

### Both CAPS- and FAP115-like regions are required for the ciliary localization and function of MpCAFA

MpCAFA consists of three distinct domains: the CAPS-like, FAP115-like, and proline-rich domains (Fig. 1). A series of constructs (Fig. S1) was prepared and expressed in Mp*cafa^ge^* plants to investigate the contribution of each domain to ciliary function and localization. CAPS-FAP115-mCitrine, which lacks the proline-rich domain, recovered its swimming speed (Fig. 10), revealing that the proline-rich domain is dispensable for MpCAFA functionality. In contrast, none of the constructs with CAPS-like or FAP115-like domains alone (CAPS, CAPS-mCitrine, FAP115, and FAP115-mCitrine), regardless of mCitrine tagging, complemented the Mp*cafa^ge^*phenotype (Fig. 10), suggesting that both regions are required for MpCAFA functionality.

**Fig. 10.**
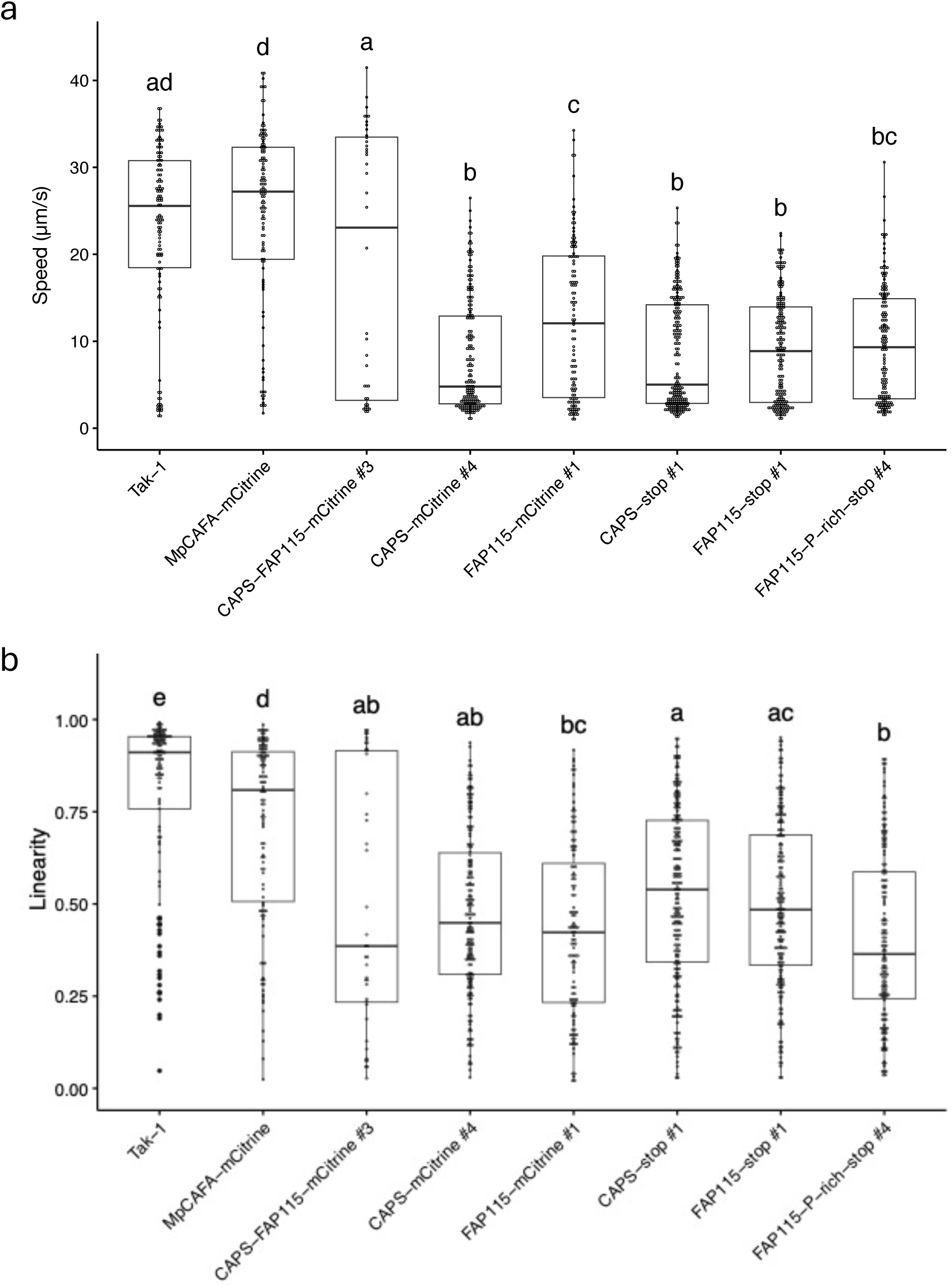
Functional analysis of each domain of MpCAFA. Box-and-whisker plots of spermatozoid swimming speed (**a**) and linearity (**b**). The upper and lower limits indicate the first and third interquartile ranges, respectively, and the middle bars indicate medians. Whiskers indicate ± 1.5× the interquartile ranges. Different lowercase letters indicate significant differences (*P* < 0.05; Tukey–Kramer test). n = 121, 114, 40, 178, 116, 216, 194, and 155 spermatozoids of Tak-1, MpCAFA-mCitrine, CAPS-FAP115-mCitrine #3, CAPS-mCitrine #4, CAPS-stop #1, FAP115-mCitrine #1, FAP115-stop #1, and FAP115-P-rich-stop #4, respectively. All the constructs were driven by the Mp*CAFA* promoter and introduced into Mp*cafa^ge^*-1 #9-2_4 plants.

Previous studies have localized mammalian CAPS (Setu et al. 2024) and algal FAP115 (Ma et al. 2019) to cilia, suggesting that the CAPS- or FAP115-like domains within MpCAFA may independently target cilia. To investigate this, we examined the subcellular localization of mCitrine-tagged proteins carrying these domains separately. Full-length MpCAFA-mCitrine and a truncated protein lacking the proline-rich domain (CAPS-FAP115-mCitrine) were consistently detected in antheridia at all developmental stages and along the entire length of the cilia (Fig. 9), indicating that the proline-rich domain is dispensable for ciliary localization. In contrast, CAPS-mCitrine (carrying only the CAPS-like domain) was detected in antheridia prior to maturation, but was absent from mature antheridia and spermatozoids (Fig. 9). FAP115-mCitrine (carrying only the FAP115-like domain) was weakly expressed in immature antheridia, localized near the periphery of the male receptacle (Fig. 9a), and was detected at the anterior end of the nucleus in spermatozoid (Fig. 9b). These findings suggest that both the CAPS- and FAP115-like domains are required for the proper ciliary localization and/or stability of MpCAFA.

## Discussion

Intracellular Ca²⁺ concentration plays a critical role in regulating ciliary movement and sperm chemotaxis in animals. However, the molecular mechanisms of Ca²⁺ signaling during sperm chemotaxis in land plants remain largely unexplored. Although the *P. patens* non-selective Ca²⁺ channel GLR has been shown to function in both sperm chemotaxis and sporophyte development (Ortiz-Ramírez et al. 2017), a comprehensive understanding of this process is lacking. In the present study, we analyzed the function of Mp*CAFA*, a gene encoding an EF-hand domain protein involved in Ca²⁺ binding, in sperm chemotaxis.

MpCAFA is unique because it contains contiguous CAPS and FAP115 domains in a single polypeptide. Although the biological significance of this structure is unknown, MpCAFA homologs are conserved among charophytes, bryophytes, pteridophytes, and gymnosperm cycads and gingko, whereas they are expressed as distinct proteins in ciliated chlorophytes and ciliates (Figs. 1 and 2; Table 1). The charophyte conjugating algae, including *Mesotaenium endlicherianum* and *Spirogloea muscicola,* have undergone secondary loss of spermatozoids, associated with the concurrent loss of CAFA. This correlation is also evident in seed plants. For example, the gymnosperm *Cycas panzhihuaensis* and *Ginkgo biloba* has both spermatozoids and CAFA, whereas angiosperms lack both (Table 1).

To elucidate the function of MpCAFA in sperm chemotaxis in *M. polymorpha*, we generated genome-edited plants by CRISPR/Cas9. Loss of Mp*CAFA* function led to a significant reduction in spermatozoid motility (Fig. 7). However, Mp*cafa*^ge^ spermatozoids retained their normal morphology, including the canonical 9 + 2 microtubule structure of the axoneme, and their chemotactic behavior and fertility. The expression of MpCAFA-mCitrine complemented the impaired spermatozoid motility in Mp*cafa*^ge^ and demonstrated a uniform distribution of MpCAFA along the entire lengths of both cilia (Fig. 9b). These results suggested that MpCAFA is a structural component of the axoneme and plays a role in the propagation of ciliary oscillations from the basal region to the distal end, but not in chemotactic behavior or fertility.

Studies on ciliary fine structures have mainly been conducted in the ciliate *Tetrahymena thermophila* and the green alga *C. reinhardtii*. Recently, the localization of microtubule inner proteins (MIPs) has been investigated using cryogenic electron microscopy (cryo-EM), revealing that MIPs interact with tubulin. RIB72, a 72-kDa ribbon-associated protein containing an EF-hand domain, binds to microtubules and localizes within the A-tubule of ciliary microtubule doublets, contributing to the ciliary beat frequency in *T. thermophila* (Stoddard et al. 2018). Interestingly, in *T. thermophila,* proteomic analysis of cilia of *rib72a/b* null mutants revealed a decrease in ciliary CAPS and FAP115 (Fabritius et al. 2021), and the mutants showed abnormal bends during the power stroke, suggesting that the structural integrity of cilia was compromised in the mutants because of the lack of these proteins (Stoddard et al. 2018). Similarly, the reduced propagation of ciliary oscillation and speed in Mp*cafa^ge^* spermatozoids can be explained by decreased ciliary rigidity because of the loss of functional MpCAFA. In *T. thermophila*, FAP115 localizes to cilia and basal bodies, and its null mutant showed decreased ciliary beat frequency and swimming speed, although cilium bending during the power stroke is not discernibly affected (Fabritius et al. 2021). In *C. reinhardtii*, RIB72 and FAP115 have been shown to localize within the A tubule (Ma et al. 2019), although their functions remain unclear. These findings demonstrate that CAPS, FAP115, and CAFA are components of the axoneme assembly and are involved in ciliary movement.

MpCAFA comprises CAPS, FAP115, and P-rich domains. To investigate the function of each domain, the truncated proteins were expressed in Mp*cafa^ge^*plants. The P-rich domain is conserved among liverworts; however, its elimination does not significantly affect the function of MpCAFA, suggesting its dispensable nature. In contrast, the CAPS and FAP115 domains are required for the proper allocation of MpCAFA to cilia. CAPS-mCitrine was abundantly present in younger antheridia, but was not detected in mature antheridia or spermatozoids, suggesting autophagic degradation during spermiogenesis. Unlike CAPS-mCitrine, FAP115-mCitrine was detected in spermatozoids; however, its localization was altered to the posterior region of the spermatozoid, where a plastid and posterior mitochondrion are present. These observations implied that the CAPS and FAP115 domains play a role in the ciliary localization and stability of MpCAFA during spermiogenesis, respectively. In *T. thermophila,* RIB72 recruits FAP115, and may also recruit CAPS, to the axoneme (Fabritius et al. 2021). *M. polymorpha* also has an RIB72 ortholog (Mp8g05900) expressed specifically in antheridia (MarpolBase Expression; Kawamura et al. 2022), but the FAP115 domain alone was not incorporated into cilia. It is not clear whether CAPS plays a role in the ciliary localization of FAP115 in animals; however, the CAPS domain may be required for the proper localization of MpCAFA in *M. polymorpha*, suggesting a similar function in animal spermiogenesis.

In this study, we identified MpCAFA as a protein involved in spermatozoid motility. Our results demonstrated that both the CAPS and FAP115 domains are indispensable for normal ciliary function in *M. polymorpha*. Despite being conserved across a wide variety of eukaryotes, CAPS and its homologs in animals have received limited attention compared with FAP115. Our findings on MpCAFA shed light on the functions of CAPS and its homologs in organisms other than *M. polymorpha*. Further investigations are required to understand the biological significance of the conserved chimeric structure of CAFA. As MpCAFA is abundant in both cilia, it can be used as a ciliary marker that does not affect spermatozoid development, swimming, or fertility.

## Acknowledgments

We thank Dr. Kazuo Yamagata (Kindai University) for recording and providing the fluorescent microscopy movie (Movie S6) used in this study. We thank Dr. Naoki Minamino (Fukuoka University), Dr. Takashi Ueda (National Institute of Basic Biology), Dr. Motomu Akita (Kindai University) for their insightful discussions. We also thank Dr. Katsura Zaitsu (Kindai University) for advice on coding R scripts. This work was supported in part by JSPS KAKENHI Grant Numbers 24112715 and 21K06237, the Japanese Association for Marine Biology (JAMBIO) (No. 35), and the Project Research of the Faculty of Biology-Oriented Science and Technology, Kindai University, 2022-2024 (No. 21-Ⅰ-2) to K.T.Y.

## Statements and Declarations

The authors declare that they have no conflict of interest.

## Supplementary data

Supplementary data are available online.

## Supplementary tables

Table S1 List of primers used in this study

Table S2 List of amino acid sequences used for phylogenetic analysis

## Legends for supplementary figures

**Fig. S1 Vector construction**

Schematic illustration of the constructs used in this study. Only the Mp*CAFA_pro_*-driven cassettes of the vectors are depicted. The backbone vectors for each construct are shown in parentheses. P, proline-rich domain; mC, mCitrine.

**Fig. S2 Mutated sequences at the target sites in female Mp*cafa^ge^* plants**

(**a**) and (**b**) Mutations introduced by CRISPR/Cas9 at the target sites 1 and 2. Nucleotide sequences of the target sites in the wild-type (WT) and female mutant (*e.g.*, 4-3) plants are shown with the translated amino acid sequence for each. Target and PAM sequences are underlined and green, respectively. The mutations are shown in red. Large deletions are indicated by the number of deleted nucleotides. Asterisks indicate the translational stops introduced during genome editing.

**Fig. S3 Morphologies of Mp*cafa^ge^* plants**

Thalli were grown from gemmae for 14 days, and the sexual organs of males (left) and females (right) are shown. Scale bar, 1 cm (Thalli).

## Legends for supplementary movies

**Movie S1 Spermatozoid motility in WT**

This movie was captured at 100 fps and is shown in real time.

**Movie S2 Spermatozoid motility in Mp*cafa*^ge^-1 #9-2**

This movie was captured at 100 fps and is shown in real time.

**Movie S3 Spermatoid motility in Mp*cafa*^ge^-2 #3-1**

This movie was captured at 100 fps and is shown in real time.

**Movie S4 Spermatozoid motility in Mp*cafa*^ge^-1 #9-2_4**

This movie was captured at 100 fps and is shown in real time.

**Movie S5 Spermatozoid motility in Mp*CAFA_pro_*:Mp*CAFA*-*mCitrine* Mp*cafa*^ge^-1 #9-2_4**

This movie was captured at 100 fps and is shown in real time.

**Movie S6 Motility of spermatozoids expressing MpCAFA-mCitrine under a fluorescent microscope**

