## Supplementary Figures for "The Mp*CAFA* gene encodes a ciliary protein required for spermatozoid motility in the liverwort *Marchantia polymorpha*"

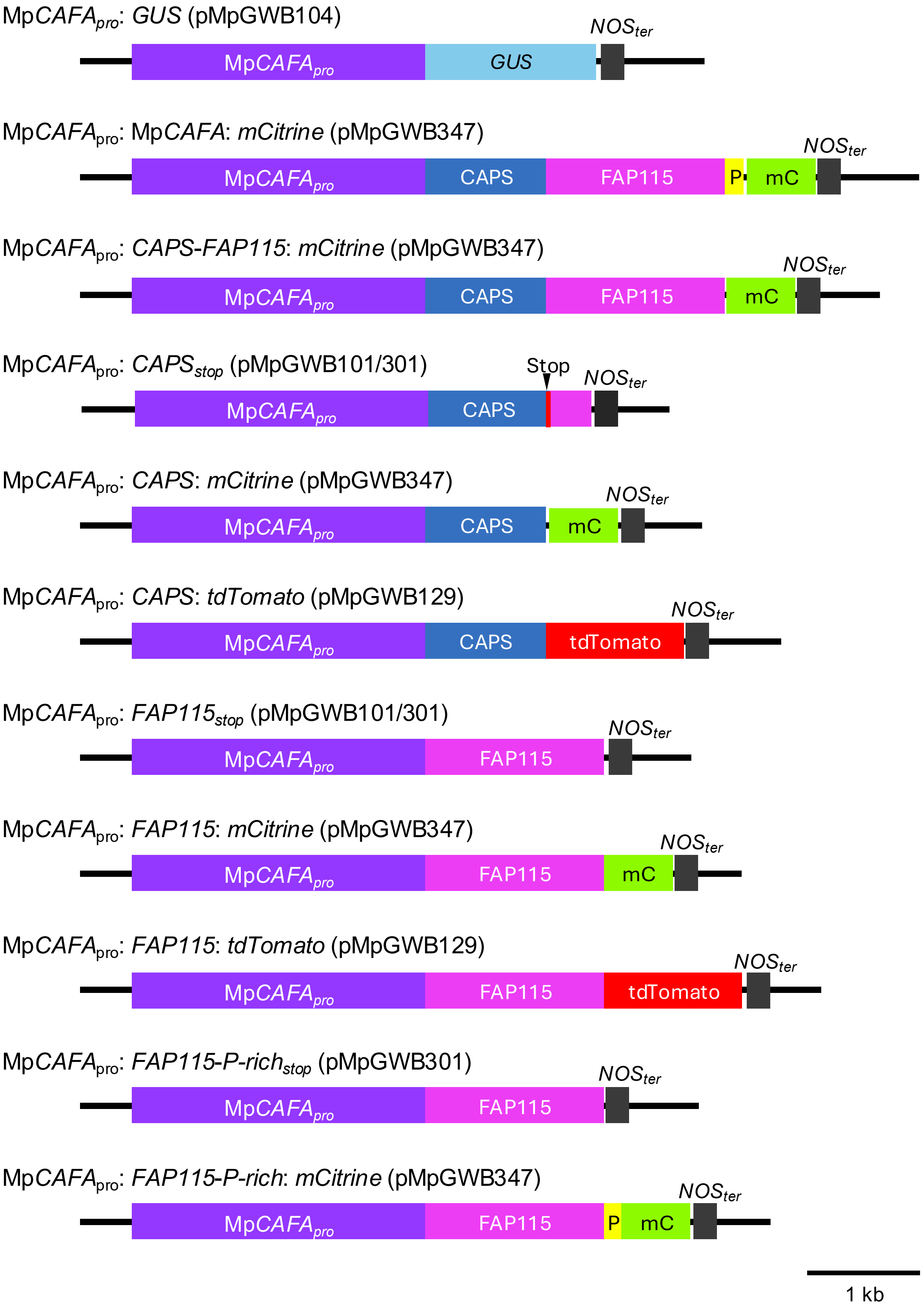


**Fig. S1 Vector construction**

Schematic illustration of the constructs used in this study. Only the Mp*CAFA*$_{pro}$-driven cassettes of the vectors are depicted. The backbone vectors for each construct are shown in parentheses. P, proline-rich domain; mC, mCitrine.

**a**

```
         M  K  T  H  C   P  D  D  K  C
WT       ATGAAGACCCACTG TCCCGATGACAAATGC

         M  K  T  H  C  S  R  *
#4-3     ATGAAGACCCACTGTTCCCGATGACAAATGC

         M  K  T  H  *
#17-1    ATGAAGACCCACTG- ----ATGACAAATGC

         M  K  T  H  L   P  D  D  L  C
#18-2    ATGAAGACCCACCT TCCCGATGACAAATGC
```

**b**

```
         L  L  H  Y    G  E  M  V  C
WT       CTGCTCCACTA  CGGCGATATGGTTTGC

         L  L  H  *
#1-2     CTGCTCCACTA  --Δ222 bp----AAG

         L  L  H  *
#13-1    CTGCTCCACTA  -GGCGATATGGTTTGC

         L  L  H  Y  G  A  G  S  K  P  Y  C  D
#22-1    CTGCTCCACTATGGAGCAGGTAGTAAACCATACTGCGAT
```

**Fig. S2 Mutated sequences at the target sites in female Mp*cafa*$^{ge}$ plants**
(**a**) and (**b**) Mutations introduced by CRISPR/Cas9 at the target sites 1 and 2. Nucleotide sequences of the target sites in the wild-type (WT) and female mutant (*e.g.*, 4-3) plants are shown with the translated amino acid sequence for each. Target and PAM sequences are underlined and green, respectively. The mutations are shown in red. Large deletions are indicated by the number of deleted nucleotides. Asterisks indicate the translational stops introduced during genome editing.

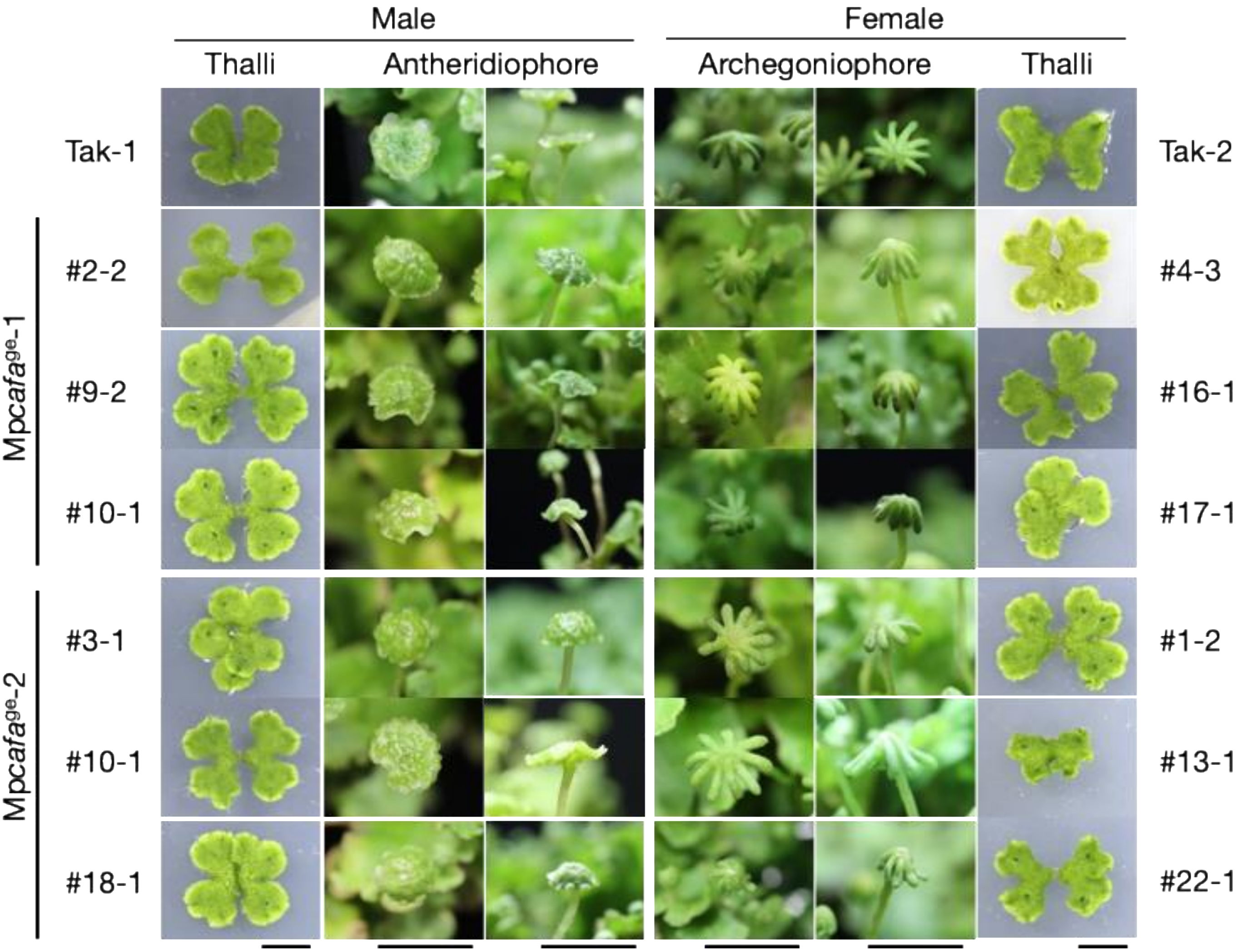


**Fig. S3 Morphologies of Mp*cafa*$^{ge}$ plants**
Thalli were grown from gemmae for 14 days, and the sexual organs of males (left) and females (right) are shown. Scale bar, 1 cm (Thalli).
